# FOLFIRINOX Combined with GPX4 Inhibition Induces Ferroptosis and Defines Redox-Based Therapeutic Subgroups in Pancreatic Cancer

**DOI:** 10.1101/2025.09.12.675778

**Authors:** Carlos Lacalle-Gonzalez, Carlos Lopez-Blazquez, Miguel Angel Hidalgo-Leon, Michael Ochienǵ Otieno, Lara Sanz-Criado, Maria Jesus Fernandez-Aceñero, Luis Ortega-Medina, Jesus Garcia-Foncillas, Javier Martinez-Useros

## Abstract

**Purpose:** Ferroptosis is a regulated form of cell death with therapeutic relevance in pancreatic ductal adenocarcinoma where Redox accumulation contributes to chemoresistance and poor prognosis. In this study, we aimed to evaluate the efficacy of combining the FOLFIRINOX regimen with a GPX4 inhibitor, RSL3, and to identify Redox-related predictive and prognosis biomarkers.

**Experimental Design:** A combination of *in vitro* and *in vivo* models were used to evaluate oxidative stress, apoptosis, ferroptosis, and the modulation of Redox proteins, GPX4 and SOD2. Furthermore, survival analyses were assessed with a 122 retrospective human pancreatic cancer cohort.

**Results:** The novel GPX4 inhibitor, RSL3, enhanced the effect of FOLFIRINOX by inducing ROS accumulation that triggered ferroptosis and apoptosis *in vitro* and a significant tumor shrinkage *in vivo* in those high ROS levels tumor cells. In contrast, SOD2 overexpression conferred resistance. Furthermore, high GPX4 and ROS expression levels were associated with shorter survival, while elevated SOD2 levels showed a subgroup with better prognosis in our cohort of 122 human pancreatic cancer cases. Thus, a novel molecular signature based on a GPX4-high/SOD2-low profile may help detect patients with the poorest clinical outcomes in pancreatic cancer.

**Conclusions:** Redox homeostasis regulates susceptibility to ferroptosis and influences treatment efficacy in pancreatic ductal adenocarcinoma. Notably, sensitivity to ferroptosis was determined not only by GPX4 levels alone, but also by the balance between ROS accumulation and SOD2-mediated antioxidant buffering. These findings support a biomarker-guided approach to treatment stratification and provide a rationale for the clinical evaluation of redox-based therapeutic strategies in pancreatic ductal adenocarcinoma.

**Translational relevance:** Ferroptosis is a non-apoptotic form of cell death with emerging therapeutic potential in pancreatic ductal adenocarcinoma. This study demonstrates that combining FOLFIRINOX with the GPX4 inhibitor RSL3 selectively induces ferroptosis in tumors with high ROS levels, both in vitro and in vivo, without increasing toxicity. Notably, sensitivity to ferroptosis is not determined by GPX4 levels alone, but rather by the tumor’s capacity to tolerate oxidative stress, shaped by ROS accumulation and compensatory SOD2 expression. In our clinical cohort, high GPX4 and elevated ROS levels correlate with poor prognosis, whereas high SOD2 expression identifies a favorable subgroup. These findings establish redox signaling as a clinically relevant variable for defining novel molecular subtypes with prognostic significance and support the use of redox biomarkers for patient stratification in pancreatic ductal adenocarcinoma. This work provides a strong rationale for the clinical evaluation of ferroptosis-inducing agents in combination with standard chemotherapy in phase I trials, guided by tumor redox profiling to enhance therapeutic precision.

## Introduction

Pancreatic ductal adenocarcinoma (PDAC) is the most prevalent form of pancreatic cancer and is the third leading cause of cancer-related mortality in both sexes. The annual number of cases closely mirrors the number of deaths, with 496,000 diagnoses and 466,000 deaths, making PDAC the deadliest type of cancer (1). Surgical resection is the primary treatment option for PDAC. Tumors located in the Vater ampulla, bile duct, or pancreatic head are typically treated with pancreatoduodenectomy (commonly referred to as the Whipple procedure), while those arising in the pancreatic body or tail require distal pancreatectomy or, in some cases, total pancreatectomy (2). Unfortunately, despite these surgical strategies, the 5-year survival rate for patients is only around 5-10% with the best adjuvant chemotherapy option available, and this figure drops near to 1% for patients without surgical options (3,4).

Currently, the best treatment option for PDAC is complete tumor resection with appropriate adjuvant chemotherapy. However, due to the late-stage diagnosis characteristic of pancreatic cancer, only 15–20% of patients qualify for this procedure. Even in cases of successful surgical resection, prognosis remains poor (5). Various strategies have been investigated to enhance outcomes in patients with resectable PDAC, and adjuvant chemotherapy has emerged as one of the most effective solutions (6). The CONKO-1 trial is the first to establish the advantages of adjuvant chemotherapy. Patients were randomized to receive either six cycles of gemcitabine at 1250 mg/m² (on days 1, 8, and 15 every 4 weeks) or observation. The study revealed a significant improvement in median progression-free survival (PFS) for the gemcitabine group (13.4 months; 95% CI: 11.4–15.3) compared to the control group (6.9 months; 95% CI: 6.1–7.8; p < 0.001), although it failed to demonstrate increased overall survival (OS) (7). Subsequent trials, such as ESPAC-4 (8) and PRODIGE-24 (9), explored alternative chemotherapy regimens to enhance patient outcomes and significantly influence the standard of care. The ESPAC-4 trial randomized resectable PDAC patients to receive gemcitabine monotherapy or a combination of gemcitabine and capecitabine for 6 months. This approach led to a 2.5-month improvement in OS. In the PRODIGE-24 trial, a modified 5-fluorouracil/leucovorin/irinotecan/oxaliplatin (FOLFIRINOX) regimen demonstrated a survival advantage of over six months compared with gemcitabine monotherapy. Consequently, FOLFIRINOX or gemcitabine/capecitabine have become the standard of care for patients with resectable PDAC. Although no head-to-head trials have directly compared these regimens, FOLFIRINOX appears to offer superior outcomes, and a comparative analysis has shown similar effectiveness (10). Recently, the NAPOLI-3 clinical trial have demonstrated that a combination therapy of liposomal irinotecan, 5-FU/leucovorin, and oxaliplatin (NALIRIFOX) outperformed gemcitabine plus nab-paclitaxel in terms of OS (11.1 months vs. 9.2 months), and PFS (7.4 months vs. 5.6 months) (11). Despite the lack of direct comparisons, it is expected that FOLFIRINOX/NALIRIFOX would be the preferred regimen by clinicians for patients with resectable and non-resectable tumors in the near future.

Despite advancements in the classic field of chemotherapy, research focused exclusively on PDAC has not led to significant breakthroughs in new oncology therapeutic areas. Targeted therapies based on immune therapy with anti-PD-1 for MSI-H/dMMR tumors or using targeted therapies against NTRK or RET when a fusion gene is present have shown some activity against PDAC (12–14). However, this evidence comes from basket studies without a clear focus on these patients, and the frequency of these alterations in routine clinical practice is very low to achieve a great benefit. One of the recent studies directed at PDAC is the phase III POLO trial, which has shown a slight survival benefit in term of PFS without an OS benefit for maintenance with olaparib in patients with germline *BRCA1/2* mutations (15). In early clinical trials, only sotorasib has emerged as a potential treatment option in phase I/II studies with *KRAS G12C* patients (16). Nevertheless, the frequency of this alteration is low (<5%), and it has not demonstrated a benefit in OS. In brief, the treatment backbone of most patients with PDAC is still a chemotherapy combination. Thus, there is a critical need for emerging therapeutic strategies that enhance chemotherapy efficacy in order to improve overall outcomes for patients with PDAC.

Ferroptosis is a regulated form of iron-dependent cell death and is characterized by lipid peroxide accumulation in cell membranes (17). Ferroptosis occurs when cellular antioxidant systems, including glutathione (GSH) and glutathione peroxidase (GPX), fail to counteract reactive oxygen species (ROS) accumulation owing to the Fenton reaction, where iron catalyzes ROS production, leading to lipid peroxidation of cellular membranes and ultimately disrupting their integrity (18). Mitochondrial metabolism, a major source of ATP production, is also a key contributor to ferroptosis through the generation of ROS (19). Most ROS are generated by the mitochondrial respiratory chain during aerobic metabolism. Ferroptosis was initially identified in cancer cells harboring *RAS* mutations, where activation of the RAS-RAF-MEK signaling pathway increases ROS production, thereby enhancing susceptibility to ferroptosis (20,21).

Due to their rapid proliferation and heightened metabolic activity, pancreatic cancer cells demand elevated energy levels, which leads to increased accumulation of ROS (22). Elevated levels of reactive oxygen species are associated with increased tumor aggressiveness and poorer prognosis in cancers such as breast cancer (23), whereas PDACs with lower ROS levels have demonstrated greater resistance to chemotherapy (24). This resistance caused by ROS is attributed to DNA damage, including double-strand breaks, which leads to the accumulation of numerous mutations. Moderate levels of ROS contribute to DNA damage and tumor progression; however, excessive ROS can trigger ferroptosis, underscoring their double-edged role in cancer biology. Targeting ROS imbalance offers a promising avenue to induce ferroptosis and eliminate cancer cells (25). High intracellular ROS levels presents a significant therapeutic advantage in PDAC by increasing cancer cell sensitivity to chemotherapy (26,27). Disrupting the balance between ROS production and detoxification systems, particularly through impairment of the GSH-GPX pathway, can lead to cellular dysfunction and ultimately induce ferroptosis (28). This approach emphasizes the therapeutic potential of leveraging iron-dependent oxidative stress to overcome chemoresistance in PDAC.

GPXs are a family of selenocysteine-containing enzymes that are critical for cellular antioxidant defense. Glutathione peroxidase 4 (GPX4) is a pivotal enzyme in the regulation of ferroptosis and acts as a guardian of cellular membrane integrity. GPX4 employs GSH as a substrate to neutralize lipid hydroperoxides, reducing them to their less reactive forms. In this process, glutathione (GSH) donates sulfhydryl groups that are oxidized to generate glutathione disulfide (GSSG), accompanied by the formation of water as a byproduct. This enzymatic activity is vital for neutralizing ROS-induced oxidative stress. If not properly controlled, this stress leads to lipid peroxidation, destabilizing cellular membranes and initiating cell death. GPX4 plays an essential role in cellular homeostasis and resistance to ferroptosis by maintaining a balance between ROS and antioxidant defense (29). GPX4 exists as a monomer with cytosolic, mitochondrial, and sperm nuclear isoforms, each of which performs specific physiological roles. Its knockout is embryonically lethal, highlighting its indispensable role in cellular homeostasis (30,31). In cancer, GPX4 has been implicated in maintaining oxidative balance, supporting epithelial-mesenchymal transition (EMT), and promoting aggressive phenotypes in PDAC (32). The enzymatic activity of GPX4 is dependent on glutathione (GSH), whose synthesis is contingent upon cysteine uptake through the system Xc⁻ antiporter. As a result, depletion of either GSH or cysteine increases the susceptibility of cancer cells to ferroptosis (33).

Inhibiting GPX4 represents a potential therapeutic strategy to eliminate PDAC cancer cells by disrupting the cellular antioxidant defense; thus, preclinical studies employing GPX4 inhibitors are urgently needed (34). Given PDAC’s aggressive nature and its resistance to standard treatments, investigating the interplay between ferroptosis and other oxidative stress modulating factors in the context of current chemotherapy regimens is critical for developing novel therapeutic approaches aimed at improving patient survival outcomes. This study aims to assess the therapeutic potential of GPX4 inhibition *in vitro* and *in vivo* as a strategy to improve the efficacy of PDAC therapy and to provide a clinical rationale for future trials.

## Material and methods

### Patient Samples

Formalin-fixed, paraffin-embedded (FFPE) tumor samples were obtained from 122 patients who underwent curative-intent duodenopancreatectomy between 2007 and 2013 at Hospital Clínico San Carlos (Madrid, Spain). All patients in this study provided written informed consent

### Tissue Microarray and Immunohistochemistry

Tissue microarrays were constructed from representative tumor regions using duplicate 1-mm cores. Immunohistochemistry (IHC) was performed on 2-3 µm FFPE sections using the Dako EnVision™ FLEX High pH kit (Agilent) following heat-induced epitope retrieval at 97 °C for 20 min in a PT-Link system, followed by overnight incubation with the following primary antibodies: anti-GPX4 (1:150, Abcam ab231174), anti-SOD2 (1:300, Abcam ab246860), and anti-8OHdG (as a surrogate ROS marker; 1:50, Abcam ab48508).

### Cell Lines

Human pancreatic cancer cell lines PL45, PANC-1, BxPC-3, and Panc04.03 were obtained from ATCC and cultured in DMEM or RPMI-1640 medium supplemented with 10% FBS and 1% P/S under standard conditions (37 °C, in a humidified atmosphere containing 5% CO₂).

### Flow Cytometry Assays

Reactive oxygen species, apoptosis, and ferroptosis were assessed by flow cytometry. Cells stained with the following reagents: carboxy-H2DCFDA (C400, Thermo Fisher), Annexin V-APC/propidium iodide (Thermo Fisher), and Mito-FerroGreen (TebuBio).

### Western Blot

The following primary antibodies were incubated overnight at 4 °C: GPX4 (Abcam ab231174, 1:1000), SOD2 (Abcam ab246860, 1:1000), and β-actin (Sigma-Aldrich A1978, 1:3000). Secondary HRP-conjugated anti-mouse (NA931V, 1:10000) and anti-rabbit (NA934V, 1:10000) antibodies were purchased from GE Healthcare.

### In vivo models

PL45 and PANC-1 cells were used to establish subcutaneous xenografts in female athymic nude mice (NU(NCr)-*Foxn1^nu^; Jackson Lab*). Treatments included: vehicle (control), FOLFIRINOX (fluorouracil 97.3 mg/kg, oxaliplatin 6.88 mg/kg, irinotecan 12.14 mg/kg, leucovorin 32.43 mg/kg) (MedChemExpress), and FOLFIRINOX + RSL3 (RAS-selective lethal 3; MedChemExpress) (5 mg/kg) as previously reported (34). Doses were extrapolated from human regimens according to FDA guidelines (35).

### Statistical Analysis

Quantitative data are presented as the mean ± 95% coefficient interval (CI). The Kolmogorov–Smirnov test was used to assess the normality of the variables. Comparisons between two parametric groups were performed using Student’s *t*-test, whereas the Mann–Whitney U test was applied for nonparametric data.

Correlations were analyzed using Pearson’s coefficient for parametric data and Spearman’s ρ for nonparametric data. Clinical associations were performed with Chi-square or Fisheŕs exact test. Survival curves based on IHC expression data were plotted using the Kaplan–Meier method and compared with the log-rank test. For mRNA analysis, TCGA, GEO, and EGA datasets were accessed via KMplot (36). Statistical analyses were conducted using IBM SPSS v26 and GraphPad Prism v8.0. Results were considered statistically significant when p-value < 0.05.

## Results

### Baseline oxidative stress predicts ferroptotic and apoptosis sensitivity to GPX4 inhibition

To explore redox biology and therapeutic sensitivity in pancreatic cancer models, we examined four human PDAC-derived cell lines (PL45, BxPC-3, PANC-1, and Panc04.03) for their baseline oxidative state and susceptibility to GPX4 inhibition by RSL3 (Fig. 1A). GPX4 expression revealed PL45 and BxPC-3 cell lines with high GPX4 expression (H-Score = 270), intermediate-to-high levels in PANC-1 cell line (H-Score = 160), and intermediate in Panc04.03 cell line (H-Score = 120) (Fig. 1B). Baseline oxidative state was measured by 8-OHdG immunostaining as a surrogate marker for ROS levels. ROS levels were undetectable in BxPC-3 and PANC-1 cell lines (H-Score = 0), low in PL45 cells (H-Score = 80), and the highest was observed in Pan04.03 cells (H-Score = 270), indicating significant heterogeneity in the oxidative status of these lines (Fig. 1B). Subsequently, all these PDAC-derived cell lines were treated with RSL3 to determine their dose-response curve. RSL3 structure features an aromatic heterocyclic core and a chloroacetamide moiety, which reacts with the selenocysteine residue in GPX4’s active site, irreversibly blocking its antioxidant activity (Fig. 1C).

**Figure 1.**
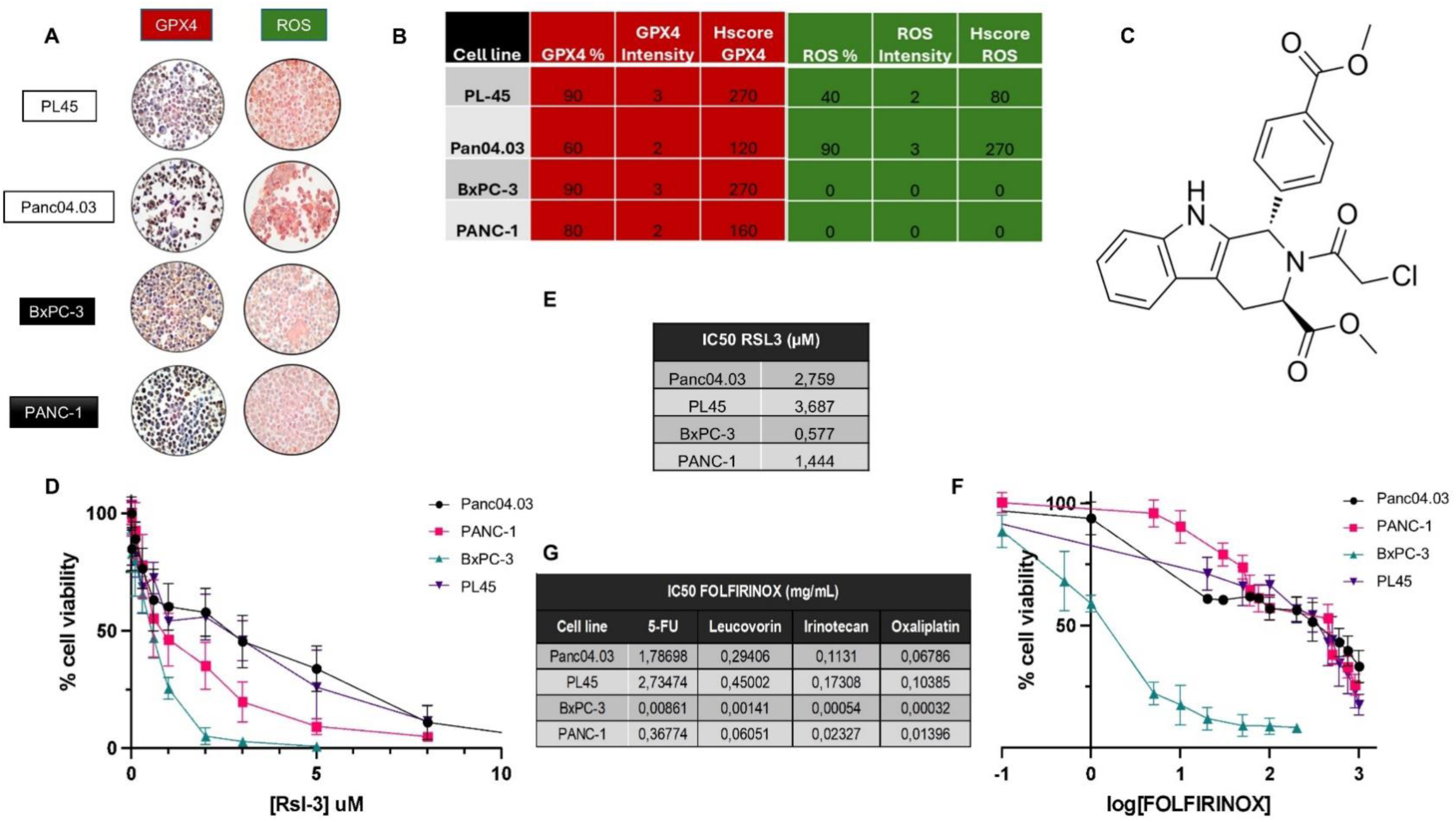
Baseline oxidative stress profiles and drug sensitivity of PDAC-derived cell lines. (A) Immunohistochemistry images for GPX4 and ROS (8-OHdG) staining in four PDAC cell lines (PL45, BxPC-3, PANC-1, and Panc04.03). (B) Semi-quantitative analysis of staining expressed as percentage of positive cells, intensity (0–3), and H-Score. (C) Chemical structure of RSL3. (D) Cell viability curves in response to RSL3 treatment (μM) in four cell lines. (E) Calculated IC₅₀ values for RSL3. (F) Cell viability curves in response to FOLFIRINOX treatment in logarithmic scale. (G) Individual IC₅₀ values for each FOLFIRINOX component (5-FU, leucovorin, irinotecan, and oxaliplatin) in mg/mL. Data represents the mean of three biological replicates.

The sensitivity to RSL3 varied considerably (Fig. 1D). The BxPC-3 cells were the most sensitive, followed by PANC-1, Panc04.03, and PL45 (Fig. 1E). One-way ANOVA with Tukey’s post hoc test revealed significant differences between BxPC-3 and PL45 cells (p = 0.0014), and between BxPC-3 and Panc04.03 cells (p = 0.0071). Interestingly, sensitivity to RSL3 did not correlate with GPX4 expression. Indeed, PL45 and BxPC-3 showed high GPX4 levels, but opposite sensitivity profiles.

FOLFIRINOX sensitivity showed distinct patterns across PDAC-derived cell lines. BxPC-3 was the most sensitive to all components, while PANC-1 showed intermediate sensitivity, and PL45 and Panc04.03 were relatively resistant. Oxaliplatin and irinotecan were generally more lethal, with BxPC-3 being the most sensitive and PL45 the most resistant. These trends largely paralleled RSL3 responses, except for Panc04.03, which was resistant to both. Full dose-response curves and R² values are provided in Supplementary Figure 1.

To determine whether baseline oxidative state influenced the effectiveness of the combination of FOLFIRINOX and RSL3, we firstly assessed ferroptosis by mitochondrial Fe2+ detection by flow cytometry (Fig. 2A–D). In PL45 cells, FOLFIRINOX alone could not increase significantly ferroptosis compared to untreated control. Remarkably, FOLFIRINOX + RSL3 significantly increased ferroptosis compared to untreated control (p = 0.0003)(Fig. 2A). In Panc04.03, a similar effect was observed, while FOLFIRINOX could not induce ferroptosis *per se*; FOLFIRINOX + RSL3 exhibited a high significance increased in ferroptosis ratio compared to untreated (p = 0.0035) (Fig. 2B). In contrast, BxPC-3 and PANC-1 showed no significant increase in ferroptosis induction by adding RSL3 to chemotherapy combination (p = n.s) (Fig 2C-D). These results confirm that PDAC-derived cell lines with the highest baseline oxidative stress are more sensitive to ferroptosis induction by RSL3 in combination with standard chemotherapy.

**Figure 2.**
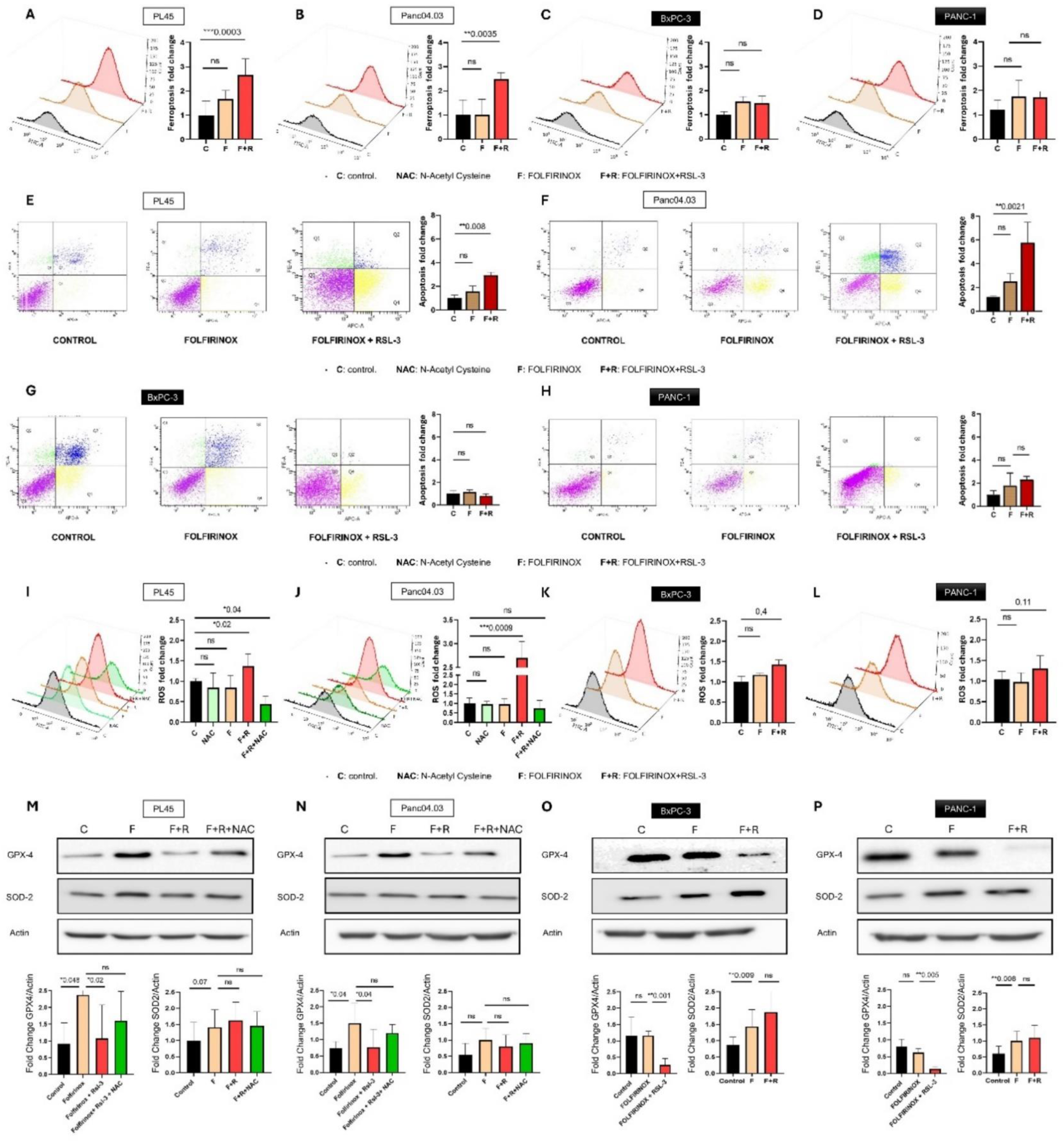
Redox response under treatment with FOLFIRINOX and RSL3 in PDAC-derived cell lines. Quantification of ferroptosis in PL45 (control mean ± 95% CI = 1.00 [0.45–1.55], FOLFIRINOX mean ± 95% CI = 1.67 [1.34–1.99]; FOLFIRINOX + RSL3 mean ± 95% CI = 2.66 [2.22–3.11]) (A), Panc04.03 (control mean ± 95% CI = 1.00 [0.52–1.48], FOLFIRINOX mean ± 95% CI = 1.01 [0.53–1.50]; FOLFIRINOX + RSL3 mean ± 95% CI = 2.48 [2.20–2.77])(B), BxPC-3 (control mean ± 95% CI = 1.00 [0.87–1.12], FOLFIRINOX mean ± 95% CI = 1.54 [1.27–1.81]; FOLFIRINOX + RSL3 mean ± 95% CI = 1.48 [1.21–1.75]) (C), and PANC-1 (control mean ± 95% CI = 1.21 [0.80–1.63], FOLFIRINOX mean ± 95% CI = 1.74 [1.13–2.34]; FOLFIRINOX + RSL3 mean ± 95% CI = 1.72 [1.46–1.97]) (D). Bar graphs show fold change in the ferroptosis ratio. Quantification of apoptosis in PL45 (control mean ± 95% CI = 1.00 [0.59–1.41], FOLFIRINOX mean ± 95% CI = 1.56 [0.81–2.31], FOLFIRINOX + RSL3 mean ± 95% CI = 2.94 [2.69–3.20]) (E), Panc04.03 (control mean ± 95% CI = 1.00 [0.66–1.34], FOLFIRINOX mean ± 95% CI = 2.51 [1.71–3.31], FOLFIRINOX + RSL3 mean ± 95% CI = 5.76 [3.02–8.50]) (F), BxPC-3 (control mean ± 95% CI = 1.00 [0.68–1.32], FOLFIRINOX mean ± 95% CI = 1.17 [0.95–1.38], FOLFIRINOX + RSL3 mean ± 95% CI = 0.82 [0.60–1.04]) (G), and PANC-1 (control mean ± 95% CI = 1.00 [0.55–1.28], FOLFIRINOX mean ± 95% CI = 1.75 [0.67–2.63], FOLFIRINOX + RSL3 mean ± 95% CI = 1.72 [0.24–2.54]) (H). Dot plots show Annexin V (x-axis) and propidium iodide (y-axis). Bar graphs show fold change in the apoptosis ratio (both early and late apoptosis). Quantification of intracellular ROS levels in PL45 (control mean ± 95% CI = 1.000 [0.914–1.086]; NAC mean ± 95% CI = 0.8460 [0.573–1.119]; FOLFIRINOX mean ± 95% CI = 0.8468 [0.625–1.069]; FOLFIRINOX + RSL3 mean ± 95% CI = 31.367 [1.132–1.601]; FOLFIRINOX + RSL3 + NAC mean ± 95% CI = 0.4424 [0.292– 0.593]) (I); Panc04.03 (control mean ± 95% CI = 1.011[0.750–1.272]; NAC mean ± 95% CI = 0.96 [0.80–1.11]; FOLFIRINOX mean ± 95% CI = 0.96 [0.67–1.25]; FOLFIRINOX + RSL3 mean ± 95% CI = 2.694 [2.328–3.061]; FOLFIRINOX +RSL3 + NAC mean ± 95% CI = 0.7553 [0.327–1.183]) (J); BxPC-3 (control mean ± 95% CI = 1.000 [0.643–1.357]; FOLFIRINOX mean ± 95% CI = 1.172 [1.084–1.267]; FOLFIRINOX + RSL3 mean ± 95% CI = 1.237 [1.170–1.305]) (K); and PANC-1 (control mean ± 95% CI = 1.013 [0.948–1.078]; FOLFIRINOX mean ± 95% CI = 0.6980 [0.768–1.185]; FOLFIRINOX + RSL3 mean ± 95% CI = 0.8678 [1.045–2.251]) (L). Bar graphs show fold change of intracellular ROS levels. Western blot quantification of GPX4 expression levels (PL45 control (mean ± 95 % CI: 0.9206 [0.1519 – 1.689]); PL45 FOLFIRINOX (mean ± 95 % CI: 2.373 [0.8821 – 3.863]); PL45 FOLFIRINOX + RSL3 (mean ± 95 % CI: 1.071 [–0.1754 – 2.318]); PL45 FOLFIRINOX + RSL3 + NAC (mean ± 95 % CI: 1.598 [0.5043 – 2.692]); Panc04.03 control (mean ± 95 % CI: 0.7333 [0.2162 – 1.250]); Panc04.03 FOLFIRINOX (mean ± 95 % CI: 1.500 [–0.01104 – 3.011]); Panc04.03 FOLFIRINOX + RSL3 (mean ± 95 % CI: 0.7667 [–0.6015 – 2.135]); Panc04.03 FOLFIRINOX + RSL3 + NAC (mean ± 95 % CI: 1.200 [0.5428 – 1.857]); BxPC-3 control (mean ± 95 % CI: 1.160 [0.4648 – 1.855]); BxPC-3 FOLFIRINOX (mean ± 95 % CI: 1.163 [0.9971 – 1.323]); BxPC-3 FOLFIRINOX + RSL3 (mean ± 95 % CI: 0.2615 [0.01392 – 0.5090]), PANC1 control (mean ± 95 % CI: 0.7926 [0.5111 – 1.074]); PANC1 FOLFIRINOX (mean ± 95 % CI: 0.6218 [0.4785 – 0.7650]); PANC1 FOLFIRINOX + RSL3 (mean ± 95 % CI: 0.1323 [0.06885 – 0.1957])); and SOD2 expression levels (PL45 control (mean ± 95% CI: 0.9972 [0.290– 1.704]); PL45 FOLFIRINOX (mean ± 95% CI: 1.409 [0.7315–2.086]); PL45 FOLFIRINOX + RSL3 (mean ± 95% CI: 1.624 [0.9175–2.330]); PL45 FOLFIRINOX + RSL3 + NAC (mean ± 95% CI: 1.458 [0.9079–2.008]); (Panc04.03 control (mean ± 95% CI: 0.55 [0.3194–1.419]); Panc04.03 FOLFIRINOX (mean ± 95% CI: 1.0 [0.1043–1.896]); Panc04.03 FOLFIRINOX + RSL3 (mean ± 95% CI: 0.8 [0.0957–1.696]); Panc04.03 FOLFIRINOX + RSL3 + NAC (mean ± 95% CI: 0.9 [0.1548–1.645]); BxPC-3 control (mean ± 95% CI: 0.861 [0.5527–1.169]); BxPC-3 FOLFIRINOX (mean ± 95% CI: 1.436 [0.7947–2.078]); BxPC-3 FOLFIRINOX + RSL3 (mean ± 95% CI: 1.873 [0.6964–4.443]); PANC1 control (mean ± 95% CI: 0.6098 [0.3236–0.896]); PANC1 FOLFIRINOX (mean ± 95% CI: 1.004 [0.6345–1.374]); PANC1 FOLFIRINOX + RSL3 (mean ± 95% CI: 1.101 [0.6139–1.588]), including representative bands and histograms normalized to actin expression for PL45 (M), Panc04.03 (N), BxPC-3 (O), and PANC-1 (P). Abbreviations: Control (C), N-acetyl-cysteine (NAC), FOLFIRINOX (F), FOLFIRINOX + RSL3 (F+R), and FOLFIRINOX + RSL3 + N-acetyl-cysteine (F+R+NAC). White boxes: RSL3-sensitive PDAC-derived cell lines; black boxes: RSL3-resistant PDAC-derived cell lines. Statistical significance was considered at p < 0.05 (n.s. = not significant).

To determine whether the enhanced cell death by ferroptosis involved apoptotic induction, we quantified apoptosis using Annexin V/PI staining and flow cytometry (Fig. 2E–H). In PL45 cells, apoptosis increased significative, especially early apoptosis, upon combination therapy with RSL3 (p = 0.008) (Fig. 2E). In Panc04.03 cell line, the effect was even more pronounced specially increasing late apoptosis (p = 0.0021) (Fig. 2F). Conversely, BxPC-3 and PANC-1 showed no significant increase in apoptosis ratio by adding RSL3 to the combination (p = n.s.) (Fig. 2G-H). Altogether, these data indicate that FOLFIRINOX + RSL3 is effective in high-ROS PDAC cell lines (PL45, Panc04.03), inducing ferroptosis and apoptosis, but not in low-ROS lines (BxPC-3, PANC-1), regardless of GPX4 expression or baseline sensitivity.

To further investigate whether the cytotoxic effect of the FOLFIRINOX + RSL3 combination was mediated by oxidative stress induction, we quantified intracellular ROS levels in four PDAC-derived cell lines using carboxy-H₂DCFDA (6-carboxy-2′,7′-dichlorodihydrofluorescein) (Fig. 2I–L). As expected, a significant increase in ROS was observed after FOLFIRINOX + RSL3 treatment exclusively in the cell lines previously identified as sensitive to the combination, PL45 and Panc04.03 (Fig. 2I–J, red bars). In contrast, cell lines resistant to FOLFIRINOX + RSL3 (BxPC-3 and PANC-1) did not exhibit ROS accumulation (Fig. 2K–L, red bars). These findings support a mechanistic link between baseline oxidative stress, and the ferroptotic response triggered by the addition of RSL3 to FOLFIRINOX.

In PL45 cells (Fig. 2I), treatment with FOLFIRINOX alone or with N-acetyl-cysteine (NAC), to reverse ROS accumulation, did not significantly alter ROS levels compared to the untreated control (all p = n.s.). Notably, co-treatment with FOLFIRINOX + RSL3 significantly enhanced ROS levels (p = 0.02). In Panc04.03 cells (Fig. 2J), the combination of FOLFIRINOX and RSL3 also induced a highly significant ROS accumulation compared to control (p = 0.0009). To determine whether the ROS increase induced by FOLFIRINOX plus RSL3 was redox-dependent, NAC was added to the combination in both cell lines (Fig. 2I-J, green bar). As hypothesized, ROS accumulation was completely reversed *in vitro* in both cell lines (p = 0.04 and p = 0.052, respectively), supporting the redox sensitivity of this effect and the role of ROS in mediating ferroptosis induced by FOLFIRINOX and RSL3.

Conversely, BxPC-3 cells (Fig. 2K) and PANC-1 cells (Fig. 2L) showed no significant changes in ROS levels under any condition (p = n.s), consistent with their lack of sensitivity to ferroptosis-inducing treatments. Collectively, these data suggest that ROS accumulation is required for RSL3-induced ferroptosis in FOLFIRINOX-treated cells, occurring only in high-ROS lines, while low-ROS lines resist ferroptosis regardless of GPX4 inhibition due to their inability to surpass a threshold of oxidative stress.

To investigate whether RSL3 modulates GPX4 levels we evaluated GPX4 by western blot in all four PDAC-derived cell lines after treatment (Fig. 2M–P). Since RSL3 is a specific GPX4 inhibitor, GPX4 levels decreased significantly upon supplementation of RSL3 to FOLFIRINOX in all four evaluated cell lines compared to FOLFIRINOX alone. We observed that NAC reversed GPX4 expression levels in both sensitive cell lines (p = n.s. vs. FOLFIRINOX). Intriguingly, GPX4 levels increased significantly in both sensitive cell lines upon FOLFIRINOX treatment (Fig. 2M–N). This indicated an adaptive redox balance to manage treatment toxicity of FOLFIRINOX. What was particularly noteworthy was that in the resistant lines BxPC-3 (Fig. 2O) and PANC-1 (Fig. 2P), FOLFIRINOX treatment did not increase GPX4 levels compared to control. These findings suggest that GPX4 induction by FOLFIRINOX represents a compensatory antioxidant response specifically in sensitive cells, rendering them vulnerable to ferroptosis upon GPX4 inhibition by RSL3. In resistant lines, the absence of GPX4 induction may explain their reduced susceptibility to ferroptosis despite effective GPX4 inhibition by RSL3. This lack of GPX4 expression in resistant lines prompted us to consider other compensatory factors that might modulate FOLFIRINOX toxicity. Consequently, we evaluated Superoxide Dismutase 2 (SOD2), which is a critical mitochondrial enzyme that protects against oxidative stress by catalyzing the dismutation of superoxide radicals into hydrogen peroxide and oxygen (37–39). The BxPC-3 cell line showed a significant increase in SOD2 levels following FOLFIRINOX and FOLFIRINOX + RSL3 treatments compared to control (Fig. 2O). Similarly, PANC-1 cells exhibited induced SOD2 expression after treatments compared to control (Fig. 2P). These results suggest that SOD2 is a primary mechanism of antioxidant adaptation in resistant PDAC-derived cell lines with low baseline oxidative stress levels. Conversely, SOD2 levels increased slightly but did not show a statistically significant increase across treatments in both sensitive cell lines compared to untreated controls (Fig. 2M-N). These data support a model in which FOLFIRINOX induces SOD2 upregulation in low-ROS, resistant cells, protecting them from ferroptosis despite GPX4 inhibition, whereas high-ROS, sensitive cells show GPX4 upregulation but not SOD2, making them more susceptible to RSL3-induced ferroptosis.

### RSL3 potentiates the effect of FOLFIRINOX *in vivo* and reduces tumor growth n high baseline oxidate stress models

To validate the translational relevance of the redox-mediated ferroptosis observed *in vitro*, we evaluated the efficacy of FOLFIRINOX and its combination with the GPX4 inhibitor RSL3 in immunodeficient murine models bearing subcutaneous xenografts. For this purpose, we selected the RSL3-sensitive PDAC-derived cell line PL45, and the RSL3-resistant PDAC-derived cell line PANC-1. Importantly, all mice (n = 36; 18 per cell line, six per arm) completed the 74-day protocol without treatment-related mortality.

In PL45 xenografts, treatment with FOLFIRINOX led to a significant reduction in tumor volume compared to the control group (mean final volume: 136.5 mm³ vs. 594.1 mm³, respectively; p = 0.0012). Notably, the addition of RSL3 to FOLFIRINOX resulted in a further and substantial tumor reduction compared to FOLFIRINOX alone (mean final volume: 19.16 mm³; p = 0.0157) (Fig. 3A). This effect was confirmed by macroscopic inspection at necropsy, where tumors in the FOLFIRINOX + RSL3 group were consistently smaller or even undetectable compared to those in the other groups (Fig. 3B). Of the twelve PL45 tumors implanted and treated with FOLFIRINOX + RSL3, eleven were evaluable: seven (63%) achieved a complete response (CR), three (27%) a partial response (PR), and one (9%) showed progressive disease (PD), yielding an objective response rate (ORR) and disease control rate (DCR) of 91%. In contrast, among the six evaluable tumors treated with FOLFIRINOX alone, two (33%) achieved CR, one (16%) had stable disease (SD), and three (50%) showed PD, resulting in an ORR of 33% and DCR of 50%. In contrast, PANC-1 xenografts exhibited minimal sensitivity to FOLFIRINOX compared to control (mean final volume: 1097 mm³ vs. 1719 mm³, respectively; p = 0.1195), and the addition of RSL3 failed to enhance therapeutic efficacy compared to FOLFIRINOX alone (mean volume: 1476 mm³; p = 0.3385) (Fig. 3C). Macroscopic assessment confirmed poor tumor control in both the FOLFIRINOX and FOLFIRINOX + RSL3 groups, with only two out of twelve tumors (16.6%) achieving complete responses (CRs) in either treatment arm, while ten out of twelve tumors (83.3%) showed progressive disease. Consequently, the objective response rate (ORR) and disease control rate (DCR) were limited to 16.6% for both regimens in this model (Fig. 3D).

**Figure 3.**
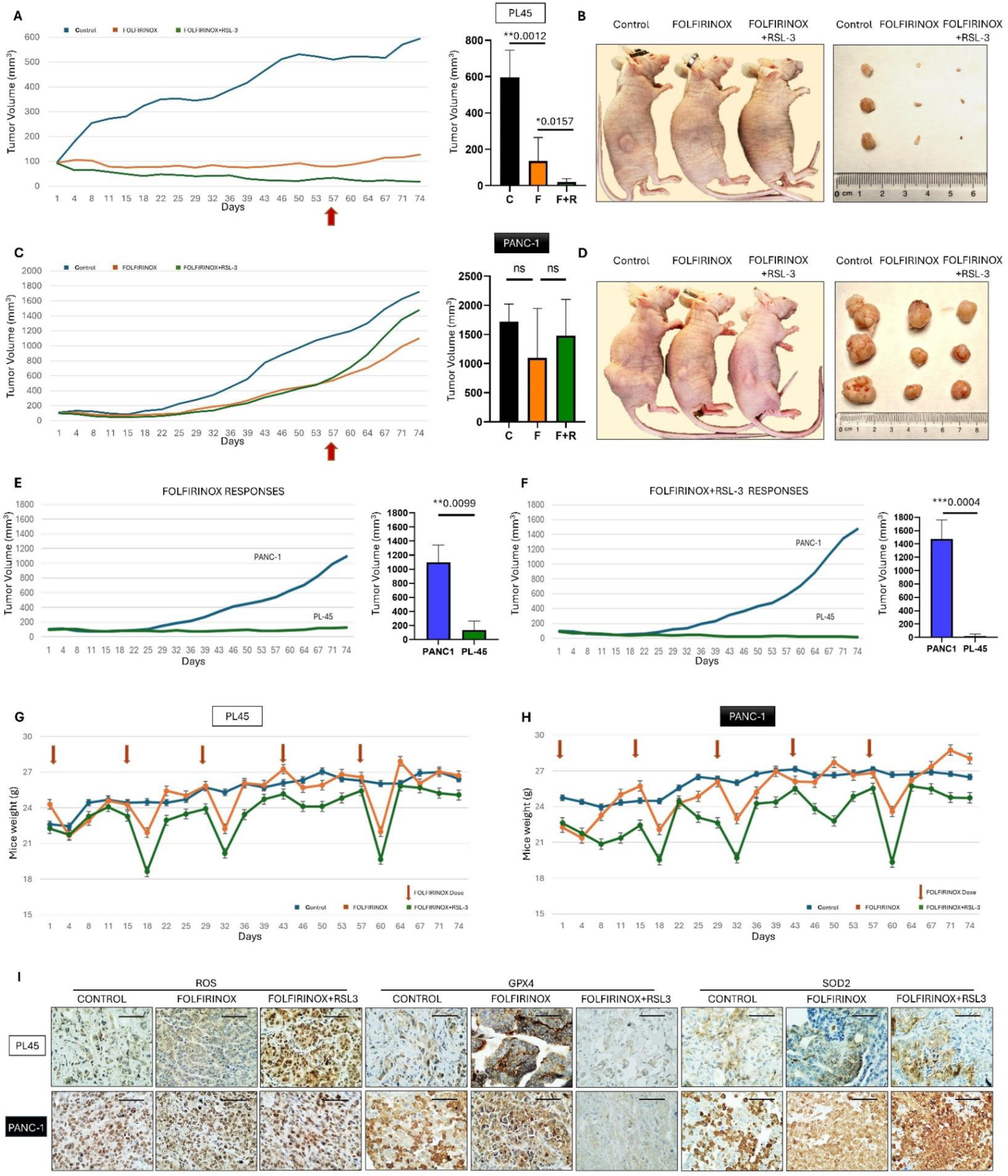
*In vivo* evaluation of FOLFIRINOX ± RSL3 in different baseline oxidative stress PDAC-derived xenograft models. (A) Tumor growth kinetics (left) and final tumor volume analysis (right) of PL45 xenografts treated with control (blue), FOLFIRINOX (orange), or FOLFIRINOX + RSL3 (green). Arrows indicate the final day of chemotherapy administration. (B) Representative photographs of mice bearing PL45 tumors (left) and excised tumors per treatment group (right). (C) Tumor growth kinetics (left) and final tumor volumes (right) of PANC-1 xenografts under identical treatment conditions. (D) Representative photographs of mice bearing PANC-1 tumors (left) and excised tumors (right). The ruler is marked in centimeters (cm). (E–F) Comparative efficacy of FOLFIRINOX monotherapy and FOLFIRINOX + RSL3 combination in PL45 and PANC-1 models, respectively. (G–H) Body weight monitoring over time for PL45 and PANC-1 xenografts, respectively. Red arrows denote FOLFIRINOX dosing days. (I) Immunohistochemical staining of xenografts for intracellular ROS, GPX4, and SOD2 expression. Scale bar represents 100 µm.

To further evaluate whether baseline oxidative stress levels of cell lines could predict responses to both standard chemotherapy and its combination with RSL3, we compared tumor volumes across both models. FOLFIRINOX alone was significantly more effective in PL45 than in PANC-1 xenografts (mean volume: 136.5 mm³ vs. 1097 mm³; p = 0.0090) (Fig. 3E). Similarly, the combination of FOLFIRINOX + RSL3 resulted in a striking contrast between PL45 and PANC-1 xenografts (mean volume: 19.16 mm³ vs. 1476 mm³; p = 0.0004) (Fig. 3F). Astonishingly, tumor regression achieved with the RSL3 combination in PL45 xenografts exceeded that observed with FOLFIRINOX alone, indicating that ferroptosis induction functions not merely as an enhancer of chemotherapy efficacy but as a potent antitumor strategy. Conversely, the lack of benefit in PANC-1 tumors treated with the same combination strongly suggests that ferroptosis sensitivity is not universally inducible. Hence, these results suggest that baseline redox status of tumor cells may influence chemotherapy efficacy, reinforcing the model-specific nature of ferroptosis sensitivity.

No signs of severe toxicity were observed throughout the treatment period. Mice treated with FOLFIRINOX or FOLFIRINOX + RSL3 exhibited mild and transient weight loss following chemotherapy cycles with no reduction exceeding 20% of baseline body weight (Fig. 3G–H). Direct observations of mice at each dosing point revealed no macroscopic differences between the FOLFIRINOX and FOLFIRINOX + RSL3 groups. Both regimens exhibited transient signs of mild toxicity compared with control animals, including decreased appetite, reduced activity, pallor, and mild fur fragility, characteristics consistent with expected chemotherapeutic stress. However, no unique adverse effects were attributable to RSL3 co-administration. Gross necropsy examination of the liver, kidneys, heart, lungs, spleen, and gastrointestinal tract revealed no treatment-related abnormalities in any group. Histopathological analysis revealed no evidence of treatment-related toxicity in any of the examined organs. In the liver, no signs of steatosis or sinusoidal obstruction syndrome were detected. Lung tissue showed no fibrosis or inflammatory changes. Renal sections displayed no tubular damage or inflammation. Cardiac tissue exhibited no myofiber necrosis or inflammatory infiltrates. The spleen showed no relevant alterations, with only occasional depletion of the white pulp. Duodenal samples showed no villous atrophy, intraepithelial lymphocytosis, or inflammation. These findings support a favorable safety profile for RSL3 at this dose level when administered in combination with FOLFIRINOX (Suppl. Fig. 2).

Subsequently, tumors were evaluated for ROS accumulation and the expression of GPX4 and SOD2, as previously assessed in *in vitro* experiments (Fig. 3I). Tumors from PL45 exhibited increased ROS expression following FOLFIRINOX treatment, which was further enhanced by the addition of RSL3. In contrast, ROS expression in PANC-1 tumors remained unchanged across treatment groups (Fig. 3I-left). As expected, both PL45 and PANC-1 xenografts exhibited decreased GPX4 expression following treatment with FOLFIRINOX + RSL3. In contrast, PL45 xenografts treated with FOLFIRINOX alone showed a slight increase in GPX4 expression compared to controls, while PANC-1 tumors remained unchanged (Fig. 3I-central). Regarding SOD2, PL45 tumors did not show increased expression in any treatment group, whereas PANC-1 tumors exhibited a slight overexpression of SOD2 in the FOLFIRINOX group, which was further elevated in the combination with RSL3 (Fig. 3I-right). These results indicate that tumors with high baseline oxidative stress are more responsive to FOLFIRINOX and RSL3, as evidenced by increased ROS levels and dynamic changes in antioxidant enzyme expression across treatments.

### Molecular subtypes based on oxidative stress modulators GPX4 and SOD2 predict clinical outcome in PDAC patients

A total of 122 patients with resectable early-stage PDAC were recruited to assess the association between survival and the expression levels of ROS, GPX4, and SOD2 in their tumor samples (Table 1). Most patients were ≥65 years of age (73%), and the sex distribution was slightly female-predominant (55.7%). Tumors were predominantly poorly differentiated (63.1%) and frequently exhibited perineural invasion (74.6%), lymph node involvement (59%), and vascular invasion (43.4%). Stage II disease was the most common (54.1%), with T2 (35.2%) and T3 (36.9%) as the most frequent T categories. Resection margins were microscopically negative (R0) in 53.3% and positive (R1) in 33.6% of cases. Adjuvant therapy was administered to 41% of patients.

**Table 1.** Clinicopathological characteristics of patients with resectable PDAC included in the study.

| Clinical characteristics | N | % | Clinical characteristics | N | % |
| --- | --- | --- | --- | --- | --- |
| Age |  |  | Grade |  |  |
| < 65 years | 25 | 20.5 | Poorly-differentiated | 77 | 63.1 |
| ≥65 years | 89 | 73 | Well-Differentiated | 38 | 31.1 |
| N/A | 8 | 6.6 | N/A | 7 | 5.7 |
| Sex |  |  | Vascular invasion |  |  |
| Male | 54 | 44.3 | No | 59 | 48.4 |
| Female | 68 | 55.7 | Yes | 53 | 43.4 |
| Adjuvant treatment |  |  | N/A | 10 | 8.2 |
| No | 57 | 46.7 | Perineural invasion |  |  |
| Yes | 50 | 41.0 | No | 21 | 17.2 |
| N/A | 15 | 12.3 | Yes | 91 | 74.6 |
| Pathologic T status |  |  | N/A | 10 | 8.2 |
| T1 | 26 | 21.3 | Stage TNM 8<sup>th</sup> edition |  |  |
| T2 | 43 | 35.2 | I | 38 | 31.1 |
| T3 | 45 | 36.9 | II | 66 | 54.1 |
| T4 | 1 | 0.8 | III | 18 | 14.8 |
| N/A | 7 | 5.7 | IV | 0 | 0 |
| Pathologic N Status |  |  | Margin status |  |  |
| N0 | 49 | 40.2 | R0 | 65 | 53.3 |
| N1 | 52 | 42.6 | R1 | 41 | 33.6 |
| N2 | 20 | 16.4 | R2 | 0 | 0 |
| N/A | 1 | 0.8 | N/A | 16 | 13.1 |
|  |  |  | TOTAL | 122 | 100 |

N: number of patients; N/A: Not available

Immunohistochemical evaluation of intratumoral ROS (Fig. 4A) revealed that high ROS levels showed a strong trend toward association with reduced PFS (median PFS: 11.1 vs. 40.1 months; *p* = 0.076) (Fig. 4B, left) and were significantly associated with shorter OS compared to tumors with low ROS levels (median OS: 18.3 vs. 48.2 months; *p* = 0.045) (Fig. 4B, right). These findings suggest that elevated oxidative stress is not only a biological hallmark of aggressive disease but also a clinically relevant prognostic factor.

**Figure 4.**
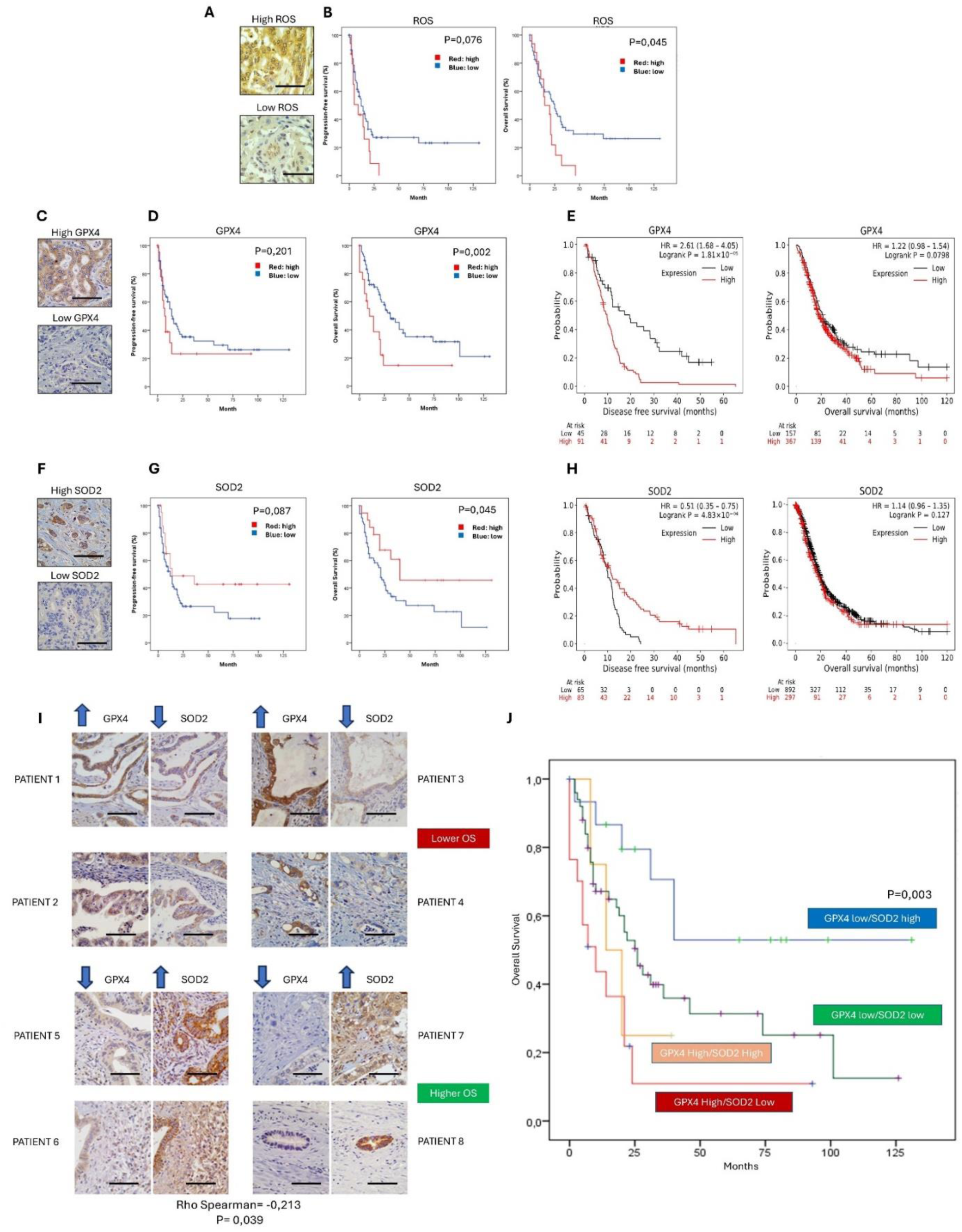
Differential expressions of ROS, GPX4 and SOD2 predict outcome in resectable PDAC patients. (A) Representative immunohistochemical staining for ROS in tumor samples showing high and low expression levels. (B) Kaplan–Meier curves of PFS (left) and OS (right) according to ROS expression levels. (C) Immunohistochemical staining for GPX4 in tumor tissues. (D) Kaplan–Meier curves for PFS (left) and OS (right) stratified by GPX4 protein expression levels. (E) Kaplan–Meier curves for disease-free survival (DFS, left) and OS (right) according to GPX4 mRNA expression levels. (F) Immunohistochemical staining for SOD2 in resected tumors. (G) Kaplan–Meier curves for PFS (left) and OS (right) stratified by SOD2 protein levels. (H) Kaplan–Meier curves for DFS (left) and OS (right) according to SOD2 mRNA expression levels. (I) Representative consecutive paired staining of GPX4 and SOD2 in tumor sections from eight patients, showing an inverse expression pattern alongside survival outcomes. (J) Kaplan–Meier curves for OS according to prognostic subgroups defined by combined GPX4/SOD2 expression. The “Redox-Aggressive” subtype (GPX4-high/SOD2-low; red) showed significantly worse survival than the “Redox-Resilient” subtype (GPX4-low/SOD2-high; blue). Scale bar = 100 μm

Similarly, elevated GPX4 protein expression (Fig. 4C) was associated with worse clinical outcomes. Patients with high GPX4 expression showed a trend toward shorter PFS (median PFS: 26.6 vs. 45.1 months; *p* = 0.201) (Fig. 4D, left) and a marked shorter OS (median OS: 22.4 vs. 53.6 months; *p* = 0.002) (Fig. 4D, right). This observation was supported by mRNA-based survival analysis, in which high GPX4 expression correlated with significantly reduced disease-free survival (DFS; HR ± 95% CI: 2.16 [1.68–4.05]; *p* = 1.81 × 10⁻⁵) (Fig. 4E, left), and showed a trend toward shorter OS (HR ± 95% CI: 1.22 [0.98–1.54]; *p* = 0.0798) (Fig. 4E, right).

In contrast, high SOD2 expression (Fig. 4F) displayed the opposite pattern, acting as a favorable prognostic factor. High SOD2 protein levels showed a strong trend toward longer PFS (median PFS: 62.0 vs. 29.9 months; *p* = 0.087) (Fig. 4G, left) and were significantly associated with prolonged OS (median OS: 72.1 vs. 40.2 months; *p* = 0.045) (Fig. 4G, right). At the mRNA level, high SOD2 expression further supported this protective effect, showing a significant association with longer DFS (HR ± 95% CI: 0.51 [0.35–0.75]; *p* = 4.83 × 10⁻⁴) (Fig. 4H, left), although no OS benefit was observed (Fig. 4H, right). Given the prognostic associations observed for ROS, GPX4, and SOD2 protein expression in overall survival analyses, we next performed a proportional hazards model including all clinicopathological variables to determine their independent contribution to patient outcomes (Suppl. Table 1). Univariate Cox analysis confirmed that age >65 years, diabetes mellitus, symptomatic presentation at diagnosis, high tumor grade, nodal involvement, high ROS, high GPX4, and low SOD2 were predictors of poorer OS (Suppl. Table 1). Multivariate analysis identified nodal involvement, high GPX4, and high ROS as independent adverse factors, whereas the protective influence of SOD2 showed a trend towards significance after adjustment (Suppl. Table 1). We then assessed the association between GPX4 and SOD2 expression levels and all clinical variables recorded in the study, but only tumor grade showed a significant association with SOD2 expression (p = 0.003) (Suppl. Table 2).

Given the opposite prognostic impact of GPX4 and SOD2, we evaluated the correlation between these two oxidative stress markers. Statistical analysis confirmed a weak but significant negative Spearman correlation (ρ = –0.213; *p* = 0.039). Moreover, opposite expression patterns were readily observed in consecutive tumor sections stained for GPX4 and SOD2. Immunohistochemistry from eight representative patients demonstrated an inverse staining pattern: high GPX4 expression coincided with low SOD2 levels, and vice versa (Fig. 4I). These findings suggest a biologically relevant antagonism between these two antioxidant systems. Therefore, survival analysis was performed using different combinations of GPX4 and SOD2 expression levels to identify an oxidative stress-related molecular signature associated with patient outcome. Interestingly, patients with a GPX4-high/SOD2-low profile experienced markedly worse outcomes, whereas those with the opposite expression pattern (GPX4-low/SOD2-high) showed prolonged survival, with the median not reached (Fig. 4J). Building on this, we defined prognostic groups according to redox-based molecular subtypes. The group characterized by GPX4-high and SOD2-low expression, termed the “Redox-Aggressive” subtype, was associated with the poorest overall survival. In contrast, patients with GPX4-low and SOD2-high expression formed the “Redox-Resilient” subtype and exhibited the most favorable prognosis. Intermediate phenotypes (GPX4-high/SOD2-high and GPX4-low/SOD2-low) showed no differences in outcomes. Survival analysis confirmed statistically significant differences in OS between all arms, underscoring the prognostic value of this dual-marker molecular signature (*p* = 0.003) (Fig. 4J). GPX4 and SOD2 define biologically distinct pancreatic cancer subtypes. High ROS and GPX4 predict poor survival, whereas high SOD2 is favorable. The inverse relationship between GPX4 and SOD2 expression refines prognostic stratification and offers a redox-based framework for tumor progression and potential therapeutic targeting.

## Discussion

Our findings reveal that ferroptosis susceptibility in PDAC depends on the balance between oxidative stress and antioxidant capacity rather than GPX4 levels per se. Ferroptosis is characterized by GPX4 inhibition and lipid peroxidation. In this context, GPX4 has been reported to act as a critical gatekeeper: cells with genetically or pharmacologically reduced GPX4 are more ferroptosis-sensitive, whereas GPX4 overexpression is broadly protective (40,41). However, our results indicate that GPX4 expression alone does not predict sensitivity to GPX4 inhibition. Instead, the magnitude of the ferroptotic response requires both impaired antioxidant defenses and a pre-existing oxidative burden, as evidenced by the lack of response in ROS-deficient cell lines such as BxPC-3 and PANC-1.

Moreover, we found that chemotherapy reprograms this redox balance in a cell type–specific manner. Cytotoxic regimens such as FOLFIRINOX induce oxidative stress, which in turn elicits adaptive antioxidant responses. For example, chemoresistant cancer cells are known to upregulate GPX4; in chemoresistant ovarian carcinoma, elevated GPX4 levels restore drug sensitivity when GPX4 is knocked down (42). Similarly, FOLFIRINOX markedly induced GPX4 expression in the ferroptosis-sensitive lines PL45 and Panc04.03. In contrast, in resistant models such as BxPC-3 and PANC-1, GPX4 induction was absent, and SOD2 was upregulated, buffering ROS levels and thereby conferring resistance to ferroptosis. This divergence reveals that not all antioxidant programs are equal: GPX4 induction facilitates RSL3 sensitivity, whereas SOD2 upregulation counteracts it. Therefore, predicting ferroptosis sensitivity requires evaluating not only baseline antioxidant levels but also the dynamic antioxidant response to therapy, as illustrated by the combination of FOLFIRINOX + RSL3.

These mechanistic insights were validated in vivo. In mouse xenografts, tumors derived from PL45 cells with a “ROS-high/GPX4-high” phenotype underwent dramatic regression when treated with FOLFIRINOX + RSL3, whereas PANC-1 tumors with a “ROS-low/GPX4-high” phenotype did not respond to the same regimen. Notably, the combination therapy was well tolerated in animals, with no overt toxicity, underscoring its potential clinical feasibility. These results indicate that the intratumoral redox signature observed in vitro persists in vivo, and that ferroptosis susceptibility can be predicted by baseline oxidative stress levels together with antioxidant response dynamics.

Importantly, results from patient samples support the clinical relevance of this model. In our PDAC cohort, tumors with high GPX4 expression and high oxidative stress had significantly worse overall survival, whereas high SOD2 expression was associated with better prognosis. Univariate Cox analysis showed that both GPX4-high and ROS-high expression conferred hazard ratios >2, whereas SOD2-high exerted a protective effect (HR: 0.472). These findings were independent of other clinicopathological factors and defined biologically distinct subtypes: GPX4-high/SOD2-low, termed “Redox-Aggressive” tumors, had poor outcomes, while GPX4-low/SOD2-high, referred to as “Redox-Resilient” tumors, had more favorable outcomes. A significant inverse correlation between GPX4 and SOD2 expression further supported the mutual exclusivity of antioxidant pathway dominance.

In summary, we propose an integrated model in which PDAC displays a broad spectrum of redox phenotypes. Tumors under high baseline oxidative stress may sustain survival through a GPX4-centered antioxidant program, whereas others rely on mitochondrial defenses such as SOD2. Chemotherapy tilts this balance: in some tumors, it shifts cells into a state where ferroptosis can be triggered by GPX4 inhibition, while in others it reinforces non-GPX4 antioxidant defenses (Fig. 5A). These insights extend the current understanding of ferroptosis biology (43) and suggest that redox-based stratification could guide personalized therapy for PDAC patients (Fig. 5B).

**Figure 5.**
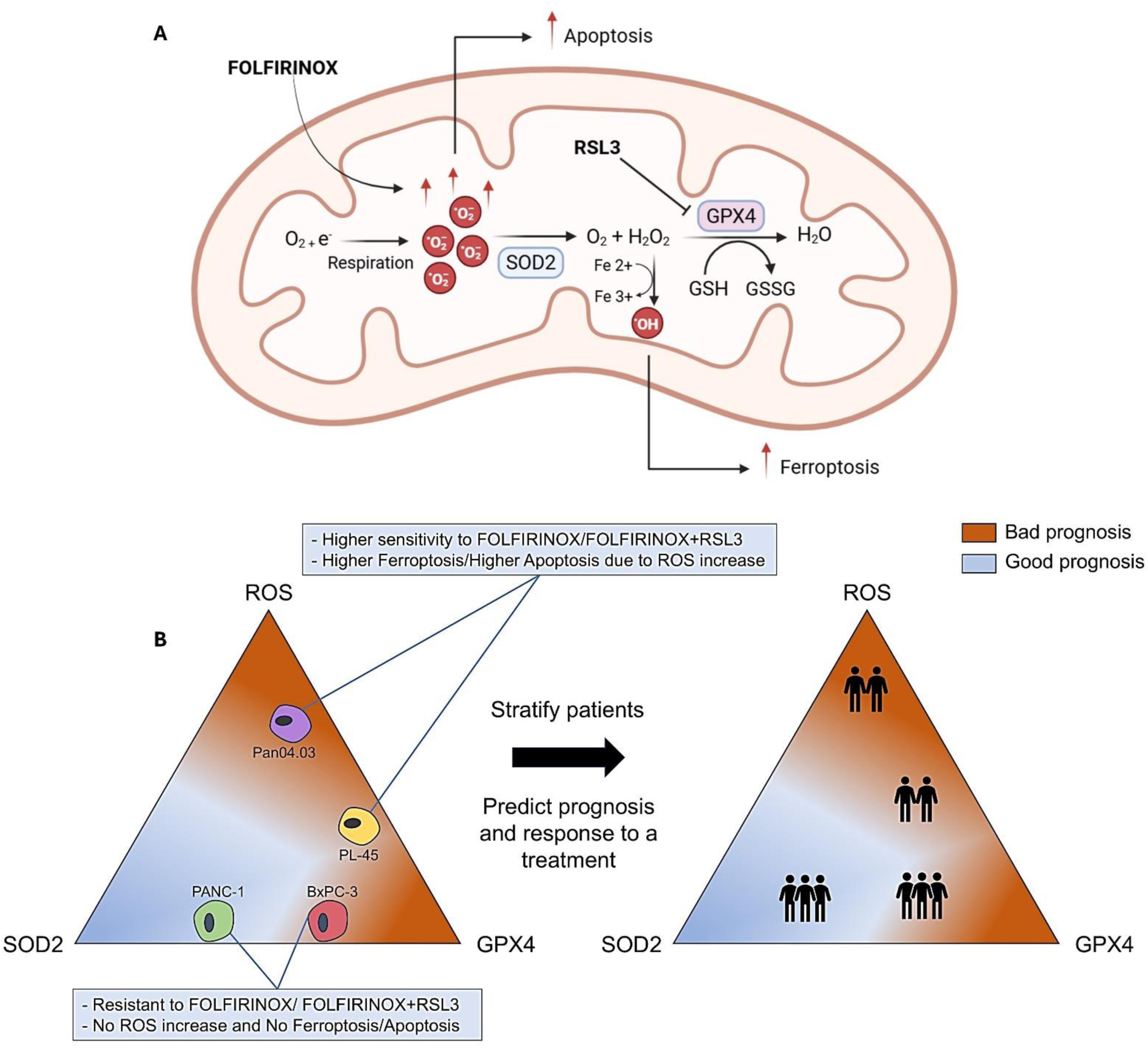
Redox-based model integrating ferroptosis sensitivity and clinical prognosis in PDAC. (A) Schematic representation of the proposed mechanism. FOLFIRINOX induces oxidative stress by increasing mitochondrial ROS. SOD2 catalyzes the dismutation of superoxide into hydrogen peroxide (H₂O₂), which is subsequently detoxified into water by GPX4. Pharmacological inhibition of GPX4 by RSL3 disrupts this antioxidant defense, leading to ROS accumulation beyond a tolerable threshold and triggering ferroptosis. The balance between ROS production and detoxification via SOD2 and GPX4 determines the intrinsic redox vulnerability of each tumor. (B) Conceptual framework linking redox biology to both therapeutic sensitivity and patient prognosis. Left panel: PDAC cell lines plotted according to baseline oxidative stress levels (ROS, GPX4, and SOD2) reveal distinct redox phenotypes that correlate with differential responses to FOLFIRINOX and its combination with RSL3. Right panel: the interplay among these three markers similarly stratifies prognosis in PDAC patients. A GPX4-high/ROS-high/SOD2-low profile identifies a high-risk, treatment-resistant subgroup, whereas a GPX4-low/ROS-low/SOD2-high phenotype is associated with improved outcomes and redox resilience. Together, this model underscores the potential of redox profiling to inform both therapeutic decision-making and patient risk stratification.

## Conclusion

This study demonstrates that redox imbalance, specifically the interplay between intracellular ROS accumulation, GPX4 induction, and SOD2 compensation, critically governs both ferroptosis susceptibility and therapeutic response in PDAC. These findings provide a strong rationale for incorporating redox biomarkers into patient stratification frameworks and support the clinical evaluation of ferroptosis-targeted therapies in molecularly selected PDAC patients.

## Authors’ Contributions

Conceptualization, C.L.-G., J.M.-U. and J.G.-F.; writing—original draft preparation, C.L.-G. and J.M.-U.; writing—review, J.M.-U.; editing, C.L.-G. and J.M.-U.; experimental procedures, C.L.-G., C.L.-B, M.A.H.-L., M.O.-O. and L.S.-C.; recruitment of patients’ samples, M.J.F.-A. and L.O.-M.; IHQ evaluation, M.J.F.-A. and L.O.-M.; visualization, C.L.-G., J.M.-U; supervision, C.L.-G., J.M.-U. and J.G.-F.; funding acquisition, J.M.-U. All authors have read and agreed to the published version of the manuscript.

## Ethics Statement

The institutional review board (IRB) of University Hospital Clinico San Carlos evaluated the present study, granting approval on 10 March 2017 with approval number n° 17/091-E. All human samples were provided by its institutional Biobank (B.0000725; PT17/0015/0040; ISCIII-FEDER).

## Data Access Statement

All relevant data are included within the paper and its supporting files. Additional data are available from the corresponding author upon reasonable request.

## Acknowledgments

The authors would like to thank Víctor Manuel Nieva and Javier Gutiérrez Corrales for their valuable assistance and technical support with the image editing software during the preparation of this manuscript. Special thanks also to Cristina Huete Cara for her kind assistance throughout the research process.

---

**Supplementary Table 1.** Uni- and multivariate Cox proportional hazard model of ROS, GPX4, SOD2 and other clinical variables for overall survival in PDAC patients.

|  | **Univariate OS (95% CI)** |  |  |  |
| --- | --- | --- | --- | --- |
|  | **HR** | **Lower** | **Upper** | **P-value** |
| Age<br>(65 years < vs > 65 years) | 2.643 | 1.340 | 5.214 | 0.005 |
| Gender<br>(Male vs Female) | 1.340 | 0.837 | 2.145 | 0.222 |
| Diabetes mellitus<br>(Yes vs. No) | 1.770 | 1.049 | 2.986 | 0.032 |
| Size<br>(< 2 cm vs > 2cm) | 1,659 | 0,921 | 2.991 | 0.092 |
| Symptoms<br>(at diagnosis) | 2.187 | 1.037 | 4.612 | 0.040 |
| Grade (low vs high) | 1.734 | 1.019 | 2.950 | 0.042 |
| Lymph nodes involved<br>(No vs Yes) | 1.899 | 1.065 | 3.386 | 0.030 |
| Vascular invasión<br>(No vs Yes) | 0.891 | 0.551 | 1.441 | 0.638 |
| Neural invasion<br>(No vs Yes) | 0.892 | 0.500 | 1.591 | 0.699 |
| GPX4 (High vs Low) | 2,473 | 1.359 | 4,503 | 0.003 |
| SOD2 (High vs Low) | 0,435 | 0,204 | 0,928 | 0,031 |
| ROS (high vs Low) | 2.151 | 1.226 | 3.775 | 0.008 |
|  | **Multivariate OS (95% CI)** |  |  |  |
| Age | 1.509 | 0.538 | 4.235 | 0.434 |
| Diabetes Mellitus | 1.668 | 0.536 | 5.189 | 0.377 |
| Symptoms | 1.040 | 0.556 | 1.948 | 0.901 |
| Grade | 0.567 | 0.223 | 1.444 | 0.234 |
| Lymph nodes involved | 3.227 | 1.135 | 9.177 | 0.028 |
| GPX4 | 5.819 | 2.050 | 16.523 | 0.001 |
| SOD2 | 0.472 | 0.154 | 1.447 | 0.187 |
| ROS | 2.987 | 1.087 | 8.206 | 0.034 |

OS: overall survival; HR: hazard ratio; CI: confidence interval; vs: versus; cm: centimetres

**Supplementary Table 2.** Association between GPX4 and SOD2 expression and clinicopathological characteristics of patients included in the study.

| Clinicopathological variables | GPX4 | SOD2 |
| --- | --- | --- |
|  | p-value | p-value |
| Age (<65 vs >65) | 0.991 | 0.776 |
| Gender (Male vs Female) | 0.286 | 0.843 |
| Diabetes Mellitus (No vs Yes) | 0.438 | 0.362 |
| Symptoms at diagnosis (No vs Yes) | 0.167* | 0.217* |
| Adjuvant treatment (Yes vs No) | 0.434 | 0.403 |
| Tumor Size ($\leq 2$ cm vs $> 2$ cm) | 0.945 | 0.104 |
| Positive Margins of resection (R0 vs R1) | 0.277 | 0.165 |
| Lymph nodes involved (No vs Yes) | 0.420 | 0.845 |
| Tumor Grade (low vs high) | 0.103 | 0.003 |
| Necrosis (No vs Yes) | 0.457 | 0.479 |
| Vascular invasion (No vs Yes) | 0.258 | 0.805 |
| Neural invasion (No vs Yes) | 0.780* | 0.623* |

777 \*Fisher's exact test was applied instead of the Chi-square test.

**Supplementary Figure 1.**
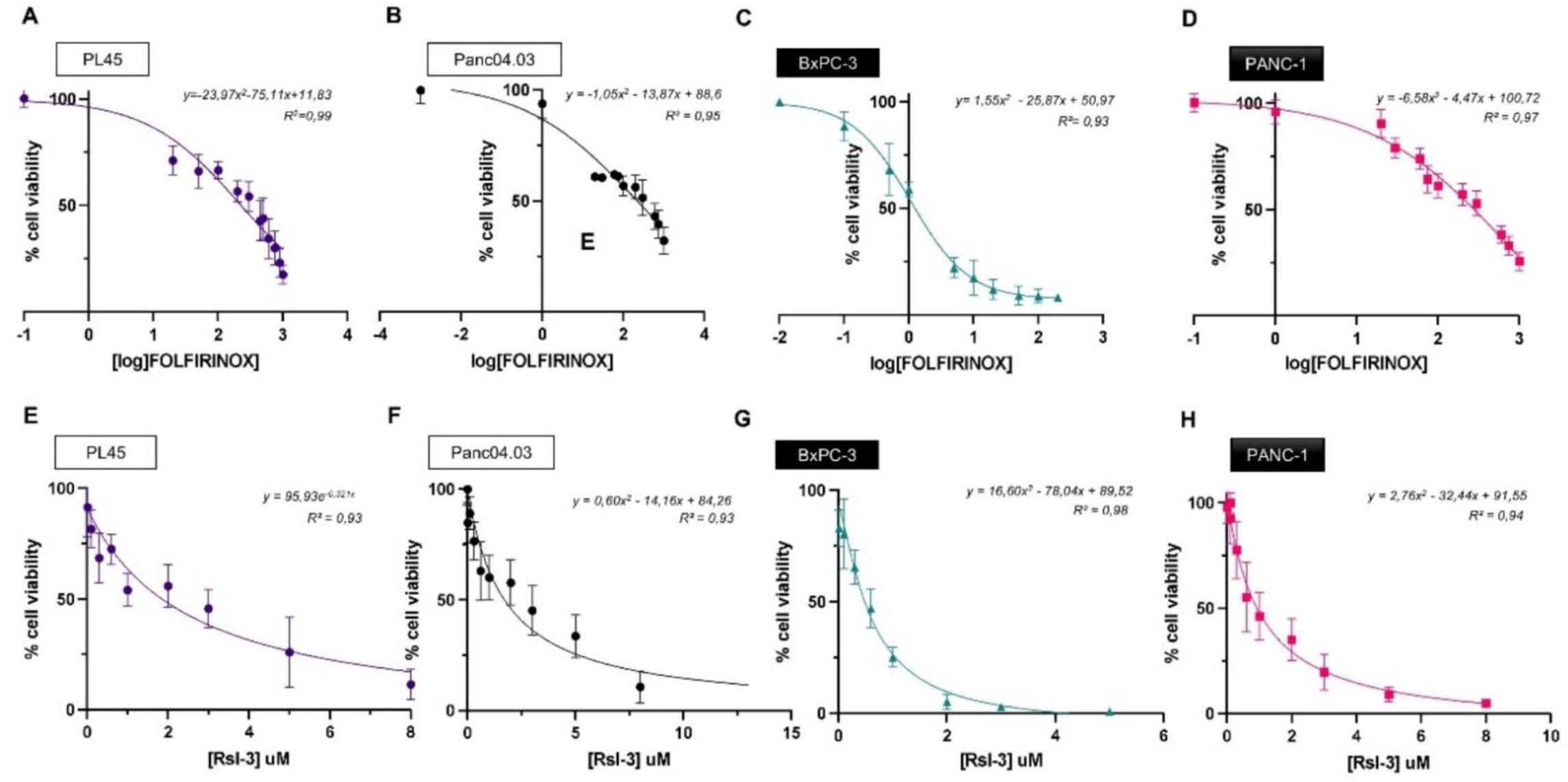
Dose-response curve for FOLFIRINOX and RSL3 in each individual PDAC-derived cell line. This figure shows dose-response curves for four pancreatic cancer cell lines (PL45, Panc04.03, BxPC-3, and PANC-1) treated with FOLFIRINOX (A-D) or RSL3 (E-H). Cell viability (%) is shown in y-axis while drug concentration is shown in x-axis. For FOLFIRINOX, the X-axis is log-transformed concentration; for RSL3, it is linear in μM. Each panel includes the best-fit equation and R² value. White boxes: RSL3-sensitive PDAC-derived cell lines; black boxes: RSL3-resistant PDAC-derived cell lines.

**Supplementary Figure 2.**
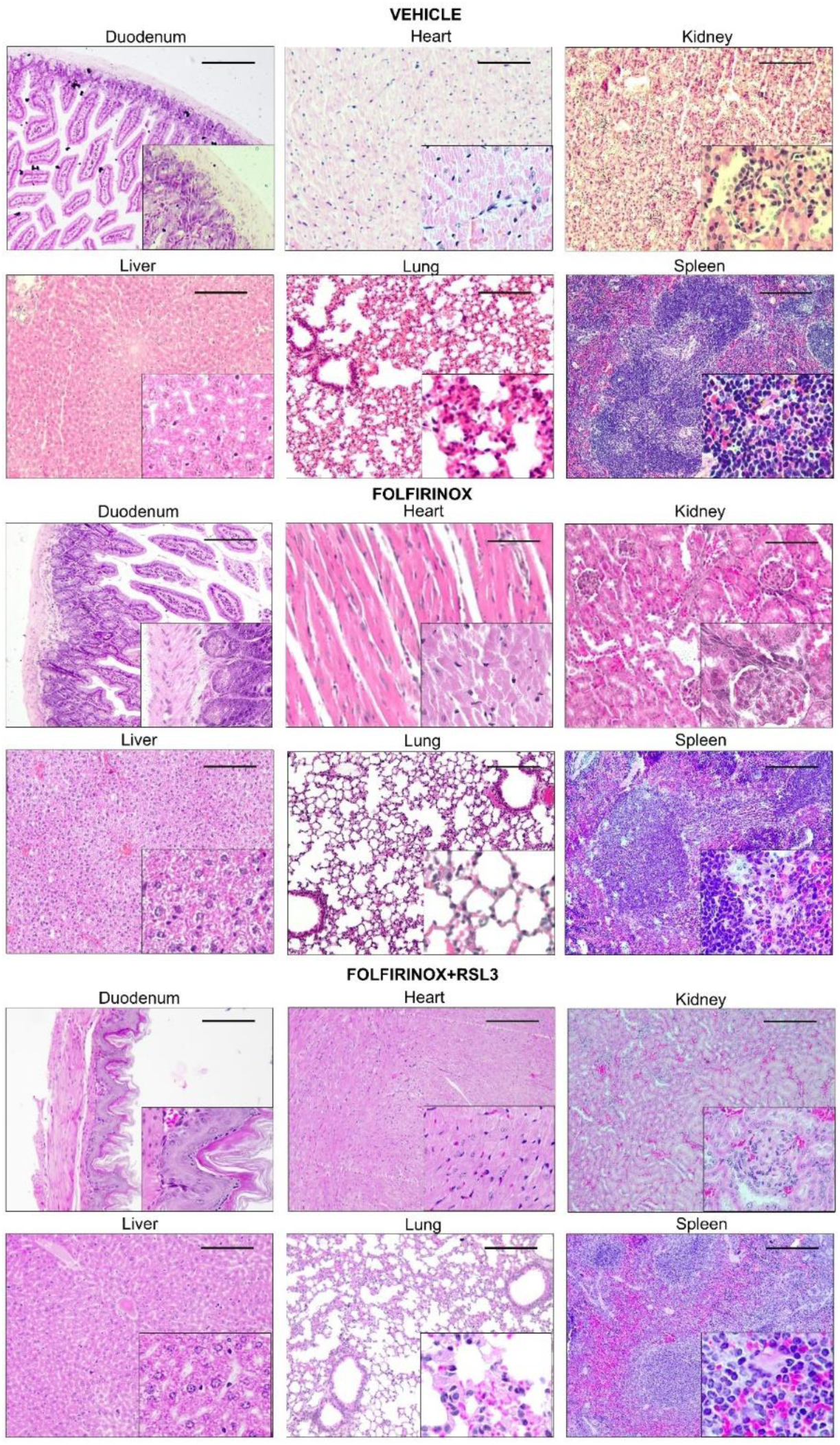
Histopathological examination of duodenum, heart, kidney, liver, lung, and spleen of mice models. This figure show representative micrographs from H&E stainings of duodenum, heart, kidney, liver, lung, and spleen of xenograft models from three experimental groups: Vehicle, FOLFIRINOX, and FOLFIRINOX + RSL3. Each organ panel includes a main image (10X) and an inset showing 60X magnification. Scale bar = 100 μm

## References

1. Siegel RL, Giaquinto AN, Jemal A. Cancer statistics, 2024. CA Cancer J Clin. 2024;74(1):12–49.

2. Conroy T, Pfeiffer P, Vilgrain V, Lamarca A, Seufferlein T, O’Reilly EM, et al. Pancreatic cancer: ESMO Clinical Practice Guideline for diagnosis, treatment and follow-up☆. Ann Oncol. 2023 Nov 1;34(11):987–1002.

3. Fan M, Deng G, Ma Y, Si H, Wang Z, Dai G. Survival outcome of different treatment sequences in patients with locally advanced and metastatic pancreatic cancer. BMC Cancer. 2024 Jan 12;24(1):67.

4. Conroy T, Castan F, Lopez A, Turpin A, Ben Abdelghani M, Wei AC, et al. Five-Year Outcomes of FOLFIRINOX vs Gemcitabine as Adjuvant Therapy for Pancreatic Cancer: A Randomized Clinical Trial. JAMA Oncol. 2022 Nov 1;8(11):1571–8.

5. Khorana AA, McKernin SE, Berlin J, Hong TS, Maitra A, Moravek C, et al. Potentially Curable Pancreatic Adenocarcinoma: ASCO Clinical Practice Guideline Update. J Clin Oncol [Internet]. 2019 Aug 10 [cited 2025 May 17]; Available from: https://ascopubs.org/doi/10.1200/JCO.19.00946

6. Conroy T, Castan F, Lopez A, Turpin A, Ben Abdelghani M, Wei AC, et al. Five-Year Outcomes of FOLFIRINOX vs Gemcitabine as Adjuvant Therapy for Pancreatic Cancer: A Randomized Clinical Trial. JAMA Oncol. 2022 Nov 1;8(11):1571–8.

7. Oettle H, Post S, Neuhaus P, Gellert K, Langrehr J, Ridwelski K, et al. Adjuvant Chemotherapy With Gemcitabine vs Observation in Patients Undergoing Curative-Intent Resection of Pancreatic CancerA Randomized Controlled Trial. JAMA. 2007 Jan 17;297(3):267–77.

8. Neoptolemos JP, Palmer DH, Ghaneh P, Valle JW, Cunningham D, Wadsley J, et al. ESPAC-4: A multicenter, international, open-label randomized controlled phase III trial of adjuvant combination chemotherapy of gemcitabine (GEM) and capecitabine (CAP) versus monotherapy gemcitabine in patients with resected pancreatic ductal adenocarcinoma: Five year follow-up. J Clin Oncol. 2020 May 20;38(15_suppl):4516–4516.

9. Conroy T, Hammel P, Hebbar M, Abdelghani MB, Wei AC, Raoul JL, et al. FOLFIRINOX or Gemcitabine as Adjuvant Therapy for Pancreatic Cancer. N Engl J Med. 2018 Dec 20;379(25):2395–406.

10. Orlandi E, Citterio C, Anselmi E, Cavanna L, Vecchia S. FOLFIRINOX or Gemcitabine Plus Nab-paclitaxel as First Line Treatment in Pancreatic Cancer: A Real-World Comparison. Cancer Diagn Progn. 2024;4(2):165–71.

11. Wainberg ZA, Melisi D, Macarulla T, Cid RP, Chandana SR, Fouchardière CDL, et al. NALIRIFOX versus nab-paclitaxel and gemcitabine in treatment-naive patients with metastatic pancreatic ductal adenocarcinoma (NAPOLI 3): a randomised, open-label, phase 3 trial. The Lancet. 2023 Oct 7;402(10409):1272–81.

12. Maio M, Ascierto PA, Manzyuk L, Motola-Kuba D, Penel N, Cassier PA, et al. Pembrolizumab in microsatellite instability high or mismatch repair deficient cancers: updated analysis from the phase II KEYNOTE-158 study. Ann Oncol Off J Eur Soc Med Oncol. 2022 Sep;33(9):929–38.

13. Berlin J, Hong DS, Deeken JF, Boni V, Oh DY, Patel JD, et al. Efficacy and safety of larotrectinib in patients with TRK fusion gastrointestinal cancer. J Clin Oncol. 2020 Feb;38(4_suppl):824–824.

14. Subbiah V, Wolf J, Konda B, Kang H, Spira A, Weiss J, et al. Tumour-agnostic efficacy and safety of selpercatinib in patients with RET fusion-positive solid tumours other than lung or thyroid tumours (LIBRETTO-001): a phase 1/2, open-label, basket trial. Lancet Oncol. 2022 Oct;23(10):1261–73.

15. Kindler HL, Hammel P, Reni M, Van Cutsem E, Macarulla T, Hall MJ, et al. Overall Survival Results From the POLO Trial: A Phase III Study of Active Maintenance Olaparib Versus Placebo for Germline BRCA-Mutated Metastatic Pancreatic Cancer. J Clin Oncol. 2022 Dec;40(34):3929–39.

16. Strickler JH, Satake H, George TJ, Yaeger R, Hollebecque A, Garrido-Laguna I, et al. Sotorasib in KRAS p.G12C–Mutated Advanced Pancreatic Cancer. N Engl J Med. 2023 Jan 4;388(1):33–43.

17. Dixon SJ, Olzmann JA. The cell biology of ferroptosis. Nat Rev Mol Cell Biol. 2024 Jun;25(6):424–42.

18. Zhou Q, Meng Y, Li D, Yao L, Le J, Liu Y, et al. Ferroptosis in cancer: from molecular mechanisms to therapeutic strategies. Signal Transduct Target Ther. 2024 Mar 8;9(1):1–30.

19. Cui K, Wang K, Huang Z. Ferroptosis and the tumor microenvironment. J Exp Clin Cancer Res. 2024 Nov 30;43(1):315.

20. Yagoda N, von Rechenberg M, Zaganjor E, Bauer AJ, Yang WS, Fridman DJ, et al. RAS–RAF–MEK-dependent oxidative cell death involving voltage-dependent anion channels. Nature. 2007 Jun;447(7146):865–9.

21. Hassannia B, Vandenabeele P, Berghe TV. Targeting Ferroptosis to Iron Out Cancer. Cancer Cell. 2019 Jun 10;35(6):830–49.

22. Pavlova NN, Zhu J, Thompson CB. The hallmarks of cancer metabolism: Still emerging. Cell Metab. 2022 Mar 1;34(3):355–77.

23. Oshi M, Gandhi S, Yan L, Tokumaru Y, Wu R, Yamada A, et al. Abundance of reactive oxygen species (ROS) is associated with tumor aggressiveness, immune response, and worse survival in breast cancer. Breast Cancer Res Treat. 2022 Jul 1;194(2):231–41.

24. Kim M, Hong WC, Kang HW, Kim JH, Lee D, Cheong JH, et al. SLC5A3 depletion promotes apoptosis by inducing mitochondrial dysfunction and mitophagy in gemcitabine-resistant pancreatic cancer cells. Cell Death Dis. 2025 Mar 7;16(1):161.

25. Moloney JN, Cotter TG. ROS signalling in the biology of cancer. Semin Cell Dev Biol. 2018 Aug 1;80:50–64.

26. Abdel Hadi N, Reyes-Castellanos G, Carrier A. Targeting Redox Metabolism in Pancreatic Cancer. Int J Mol Sci. 2021 Feb 3;22(4):1534.

27. Durand N, Storz P. Targeting reactive oxygen species in development and progression of pancreatic cancer. Expert Rev Anticancer Ther. 2017 Jan;17(1):19–31.

28. Jagust P, Alcalá S, Jr BS, Heeschen C, Sancho P. Glutathione metabolism is essential for self-renewal and chemoresistance of pancreatic cancer stem cells. World J Stem Cells. 2020 Nov 26;12(11):1410–28.

29. Yu Y, Yan Y, Niu F, Wang Y, Chen X, Su G, et al. Ferroptosis: a cell death connecting oxidative stress, inflammation and cardiovascular diseases. Cell Death Discov. 2021 Jul 26;7(1):193.

30. Ufer C, Wang CC. The roles of glutathione peroxidases during embryo development. Front Mol Neurosci. 2011 Jul 28;4:11531.

31. Berndt C, Alborzinia H, Amen VS, Ayton S, Barayeu U, Bartelt A, et al. Ferroptosis in health and disease. Redox Biol. 2024 May 30;75:103211.

32. Peng G, Tang Z, Xiang Y, Chen W. Glutathione peroxidase 4 maintains a stemness phenotype, oxidative homeostasis and regulates biological processes in Panc-1 cancer stem-like cells. Oncol Rep. 2019 Feb;41(2):1264–74.

33. Badgley MA, Kremer DM, Maurer HC, DelGiorno KE, Lee HJ, Purohit V, et al. Cysteine depletion induces pancreatic tumor ferroptosis in mice. Science. 2020 Apr 3;368(6486):85–9.

34. Yang J, Mo J, Dai J, Ye C, Cen W, Zheng X, et al. Cetuximab promotes RSL3-induced ferroptosis by suppressing the Nrf2/HO-1 signalling pathway in KRAS mutant colorectal cancer. Cell Death Dis. 2021 Nov 13;12(11):1079.

35. Research C for DE and. Estimating the Maximum Safe Starting Dose in Initial Clinical Trials for Therapeutics in Adult Healthy Volunteers [Internet]. FDA; 2018 [cited 2025 May 18]. Available from: https://www.fda.gov/regulatory-information/search-fda-guidance-documents/estimating-maximum-safe-starting-dose-initial-clinical-trials-therapeutics-adult-healthy-volunteers

36. Posta M, Győrffy B. Analysis of a large cohort of pancreatic cancer transcriptomic profiles to reveal the strongest prognostic factors. Clin Transl Sci. 2023;16(8):1479–91.

37. Vrzgula M, Mihalik J, Teslik J, Hodorova I. Differential expression of various isoforms of superoxide dismutase in the cells of the human exocrine pancreas. Bratisl Lek Listy. 2024;125(9):539–43.

38. Peoples JN, Saraf A, Ghazal N, Pham TT, Kwong JQ. Mitochondrial dysfunction and oxidative stress in heart disease. Exp Mol Med. 2019 Dec;51(12):1–13.

39. Palma FR, He C, Danes JM, Paviani V, Coelho DR, Gantner BN, et al. Mitochondrial Superoxide Dismutase: What the Established, the Intriguing, and the Novel Reveal About a Key Cellular Redox Switch. Antioxid Redox Signal. 2020 Apr 1;32(10):701–14.

40. Li J, Cao F, Yin H liang, Huang Z jian, Lin Z tao, Mao N, et al. Ferroptosis: past, present and future. Cell Death Dis. 2020 Feb 3;11(2):1–13.

41. Sui X, Zhang R, Liu S, Duan T, Zhai L, Zhang M, et al. RSL3 Drives Ferroptosis Through GPX4 Inactivation and ROS Production in Colorectal Cancer. Front Pharmacol [Internet]. 2018 Nov 22 [cited 2025 May 21];9. Available from: https://www.frontiersin.org/journals/pharmacology/articles/10.3389/fphar.2018.01371/full

42. Feng Q, Hao S, Fang P, Zhang P, Sheng X. Role of GPX4 inhibition-mediated ferroptosis in the chemoresistance of ovarian cancer to Taxol in vitro. Mol Biol Rep. 2023 Dec;50(12):10189–98.

43. An X, Yu W, Liu J, Tang D, Yang L, Chen X. Oxidative cell death in cancer: mechanisms and therapeutic opportunities. Cell Death Dis. 2024 Aug 1;15(8):1–20.

