## Supplementary Materials & Methods for "FOLFIRINOX Combined with GPX4 Inhibition Induces Ferroptosis and Defines Redox-Based Therapeutic Subgroups in Pancreatic Cancer": Supplemtary material M&M.pdf

### Material and methods

#### Patient Samples

Formalin-fixed, paraffin-embedded (FFPE) tumor samples were obtained from 122 patients who underwent curative-intent duodenopancreatectomy between 2007 and 2013 at Hospital Clínico San Carlos (Madrid, Spain). Samples were provided by the Institutional Biobank (B.0000725; PT17/0015/0040; ISCIII-FEDER) after approval by the Institutional Ethics Committee (no. 17/091-E, March 10, 2017). All patients had a confirmed histopathological diagnosis of pancreatic ductal adenocarcinoma, along with a comprehensive clinicopathological dataset.

#### Tissue Microarray and Immunohistochemistry

Tissue microarrays were constructed from representative tumor regions using duplicate 1-mm cores. Immunohistochemistry (IHC) was performed on 2-3 µm FFPE sections using the Dako EnVision™ FLEX High pH kit (Agilent) following heat-induced epitope retrieval at 97 °C for 20 min in a PT-Link system. Endogenous peroxidase was blocked using EnVision FLEX peroxidase blocking reagent (Agilent), followed by overnight incubation with the following primary antibodies: anti-GPX4 (1:150, Abcam ab231174), anti-SOD2 (1:300, Abcam ab246860), and anti-8OHdG (as a surrogate ROS marker; 1:50, Abcam ab48508). Staining was visualized with 3,3′-diaminobenzidine (DAB, Agilent) and counterstained with Harris hematoxylin (Sigma-Aldrich). Positive controls were selected based on the data from The Human Protein Atlas (www.proteinatlas.org, accessed on Jun 2024). Immunoreactivity was independently evaluated by two pathologists (M.J.F.-A. and L.O.-M.) by using the H-score method. The H-score was calculated by multiplying the percentage of positively stained cells (0–100%) by the staining intensity (1 = weak, 2 = moderate, 3 = strong), resulting in a score ranging from 0 to 300 as follows:

$$H - Score = (1 \times \%_{Low}) + (2\ \times \%_{Medium}) + \left(3\ \times \%_{High}\right)$$

Cell Lines

Human pancreatic cancer cell lines PL45, PANC-1, BxPC-3, and Panc04.03 were obtained from ATCC and cultured in DMEM or RPMI-1640 medium supplemented with 10% fetal bovine serum and 1% penicillin/streptomycin under standard conditions (37 °C, in a humidified atmosphere containing 5% $CO_2$). Mycoplasma testing was routinely performed (Lonza MycoPCR Detection Kit).

Flow Cytometry Assays

Reactive oxygen species (ROS), apoptosis, and ferroptosis were assessed by flow cytometry. Cells were harvested after 24 hours of treatment and stained with the following reagents: carboxy-H2DCFDA (C400, Thermo Fisher), Annexin V-APC/propidium iodide (Thermo Fisher), and Mito-FerroGreen (TebuBio). Samples were analyzed using a BD FACSCanto II cytometer and FACSDiva software version 6.1.3. ROS and ferroptosis were measured in the FITC (fluorescein isothiocyanate) channel, apoptosis was detected in the APC (allophycocyanin) channel (Annexin V-APC), and propidium iodide was detected in the PE (phycoerythrin) channel.

Western Blot

Cells were lysed in RIPA buffer and protein extract was quantified with the commercial kit Pierce™ BCA Protein Assay Kit (Thermo Fisher-Scientific). SDS-

PAGE gels were loaded with 30 µg of total protein per sample and transferred to PVDF membranes. The following primary antibodies were incubated overnight at 4 °C: GPX4 (Abcam ab231174, 1:1000), SOD2 (Abcam ab246860, 1:1000), and β-actin (Sigma-Aldrich A1978, 1:3000). Secondary HRP-conjugated anti-mouse (NA931V, 1:10000) and anti-rabbit (NA934V, 1:10000) antibodies were purchased from GE Healthcare. The signal was detected using enhanced chemoluminescence (ECL Prime; Amersham Pharma Biotech) in the Amersham Imager 600 (GE Healthcare).

<u>In vivo models</u>

PL45 and PANC-1 cells were used to establish subcutaneous xenografts in female athymic nude mice (NU(NCr)-*Foxn1*$^{nu}$*; Jackson Lab*) (six per group, two tumors per mouse, one per flank). Cells were injected in RPMI:Matrigel (1:1) and allowed to grow until the tumors reached 100 mm$^3$. Treatments included: vehicle (control), FOLFIRINOX (fluorouracil 97.3 mg/kg, oxaliplatin 6.88 mg/kg, irinotecan 12.14 mg/kg, leucovorin 32.43 mg/kg)(MedChemExpress), and FOLFIRINOX plus RSL3 (RAS-selective lethal 3; MedChemExpress)(5 mg/kg) as previously reported (37). Doses were extrapolated from human regimens according to FDA guidelines (38). Mice received intraperitoneal administration of FOLFIRINOX every 14 days, and RSL3 was co-administered with FOLFIRINOX intraperitoneally on days 1, 4, 8, and 11 of the 14-day cycle. Drug dilutions were prepared according to compound-specific solubility requirements: fluorouracil and oxaliplatin were dissolved in PBS, whereas irinotecan, leucovorin, and RSL3 were diluted in a vehicle consisting of 10% DMSO, 40% PEG-300, 5% Tween-80, and 45% saline. Xenograft tumour volume was measured twice per week. Toxicity was assessed based on body weight, clinical observations, necropsy findings, and histopathological examination of the duodenum, liver, heart, lungs, spleen, and kidneys. DietGel® Recovery (ClearH2O) was provided as nutritional support to all mice receiving chemotherapy.

Statistical Analysis

Quantitative data are presented as the mean ± standard deviation (SD). The Kolmogorov–Smirnov test was used to assess the normality of the variables. Comparisons between two parametric groups were performed using Student's *t*-test (paired or unpaired, as appropriate), whereas the Mann–Whitney U test was applied for nonparametric data. Correlations were analyzed using Pearson's coefficient for parametric data and Spearman's ρ for nonparametric data.

Progression-free survival (PFS) was defined as the time from surgery to recurrence, and overall survival (OS) as the time from surgery to death. Survival curves based on IHC expression data were plotted using the Kaplan–Meier method and compared with the log-rank test. The univariate Cox proportional hazards model was employed to estimate hazard ratios and 95% confidence intervals for ROS, GPX4, SOD2, and other clinicopathological variables. Only variables with statistical significance in the univariate analysis were included in the multivariate Cox regression model. As the ROC curve did not provide an appropriate cutoff point, H-score thresholds were set at the 75th percentile (p75) to stratify patients into high- and low-expression groups. For validation, survival analyses were performed using publicly available independent mRNA datasets (TCGA, GEO, and EGA) accessed via KMplot (39). Statistical analyses were conducted using IBM SPSS v26 and GraphPad Prism v8.0. Results were considered statistically significant when p-value < 0.05.
